# Refined novel techniques for long term cryo-storage using vitrification and laser warming

**DOI:** 10.1101/2023.01.04.521297

**Authors:** Chiahsin Lin, Wen-Chung Hsieh, Kanokpron Loeslakwiboon, Cheng-Liang Huang, Ting-Chun Chen, Sujune Tsai

## Abstract

Vitrification and ultra-rapid laser warming technique is an important approach for cryopreservation of animal embryos and oocytes, as well as other cells with medicinal, genetic, and agricultural value. We believe that the long term cryo-storage after vitrification and for the following laser warming has only been achieved to date using our customized device which was first attempted to be developed in our study. In the present study, we focused on developing alignment and bonding techniques for special cryo-jig which were assembling jig tool and jig holder in one piece. This newly produced customized cryo-jig was demonstrated to have significantly high laser striking accuracy of 95% and a successful rewarming rate of 62%. This study was experimentally demonstrate an refined novel device for improvement of laser striking accuracy after long term cryo-storage using vitrification and laser warming technique. In addition, the customized device described herein was successfully applied to a biological sample with over a thousand coral larvae in long term cryo-storage (The first cryo-repository for coral larvae; the results related to coral cryobanking and repositry described in this preprint will be published separately in details). We anticipate that our core findings will provide further examples of cryobanking applications that use vitrification and nano-laser warming to help a wide range of cells and tissues from diverse species.

## INTRODUCTION

Cryopreservation is widely utilized for long-term storage, despite the fact that freeze-thawing methods can severely decrease gamete function. The cryopreservation technique is defined by the balance between freezing rate and the need to accomplish regulated cellular dehydration and minimum intracellular ice formation (Leibo et al., 1981). Over the last few decades, cryopreservation techniques have become a well-established approach of preserving cells and tissues. This advancement has had a substantial influence on a variety of fields, including humans (Parmegiani et al., 2011), cows (Hwang and Hochi, 2014) and mice (Jin and Mazur, 2015). It is clear that cryopreservation of mammalian embryos and oocytes can currently be carried out by slow cooling or vitrification. While there are significant differences between these two cryopreservation methods, both require control and optimization of conditions during each stage of the procedure. In addition, cryopreservation of gametes, embryos, and embryonic cells has become a tremendous value in aquatic biotechnologies, which become a successful approach for propagating economically important species, protect endangered species, and maintain genetic diversity (Tsai and Lin, 2012). To date, the spermatozoa of more than 300 species of fish have been cryopreserved; however, research on the cryopreservation of fish oocytes and embryos remains challenging and is still in its initial stages.

During the past decade, vitrification has been acknowledged as a safe and promising alternative to conventional slow-rate freezing (Tsai et al., 2015). Vitrification can supercool biomaterials from vertebrate and invertebrate at ultra-fast cooling rate to avoid ice crystal formation by using high concentrations of cryoprotectant as one of ways and rapid cooling rates to transition directly from a solution to a clear, noncrystalline, and solid-state resembling glass (Tsai et al., 2015, 2016). In recent years, another encouraging approach for cryopreservation of germplasms of vertebrate species is the development of laser-warming with gold nanoparticles techniques to effectively reanimate vitrified zebrafish embryos (Khosla et al., 2017), and mouse oocytes/embryos (Jin et al., 2014; Jin and Mazur, 2015) that were able to resume lives after vitrification and laser warming. Laser nanowarming can aid in the rescue of cryoprotectant-sensitive coral larvae that are unable to be effectively cooled, as well as provide rapid and uniform warming rates for coral larvae to avoid further damage by ice crystallization during laser warming (Daly et al., 2018). Other notable achievements include the first successful cryopreservation of coral larvae with the presence of Symbiodiniaceae in larvae from *Seriatopora caliendrum* and *Pocillopora verrucosa* that have vertical transmission of algal symbionts were able to settle after vitrification and laser warming (Cirino et al., 2019, 2021). Furthermore, the long term cryo-storge after vitrification and for the following laser warming can only be achieved using our customized device which was first attempted to be developed in our prior study (Narida et al., 2022), however, this device required two jigs fitting positions and assembly Cryotop procedures to conduct high laser striking accuracy and effective rewarming. It is an urgent need to improve the efficiency and sustainability of innovative solutions for the future long-term cryobanking.

Thus according recent work from our laboratory has shown that two types of cryo-jig (cryo-jig A and B) were utilized to load and warm the sample for the vitrification and laser warming method, and the cryo-jig had a laser striking accuracy of 76% (Narida et al., 2022). It might be caused by imperfect alignment of laser machine with the jig axis circle, which was not centered on the origin. The goal of this research is to optimize the mechanical arrangement of the customized device and cryo-jig in order to allow the laser beam to strike a designated target and achieve greater laser striking accuracy and successful laser-rewarming without recrystallization.

## MATERIALS AND METHODS

### Alignment between customized device and laser machine

The precision alignment of the laser machine was essential for the laser to strike the sample (Fig 1). The laser machine (LaserStar Technologies Corporation, Riverside, USA) was equipped with a stereomicroscope with an ocular reticle, which facilitated sample observation and alignment within the laser beam. The specially self-customized cryo-jig was used to lower the sample into liquid nitrogen for vitrification, raise the sample into the laser beam focus zone, and trigger the laser for warming. The modified version of the cryo-jig allowed easy replacement of the cryo-stick blade after each laser pulse to ensure consistent and reproducible sample warming. Laser beam was tested by firing the beam through the laser target site. The laser beam directly passed through the center of the reticule without hitting reticle margin in order to determine the device position relatives to laser machine. A drop of vitrification solution (VS; 0.8ul) was placed on the self-made cryo-stick, which was centered in the reticles of the cryo-jig. The cryo-jig was secured steady at a specifically designated location and a laser beam was emitted again to ensure perfect alignment and that the drop was struck. The energy provided by a single pulse can be varied by changing the input voltage and pulse period. A laser calibration system was generated using a laser power meter to determine the energy in a laser pulse. The alignment process was repeated at least every ten samples in order to ensure the laser pulse footprint fully covered the sample each time successfully.

**Fig 1.**
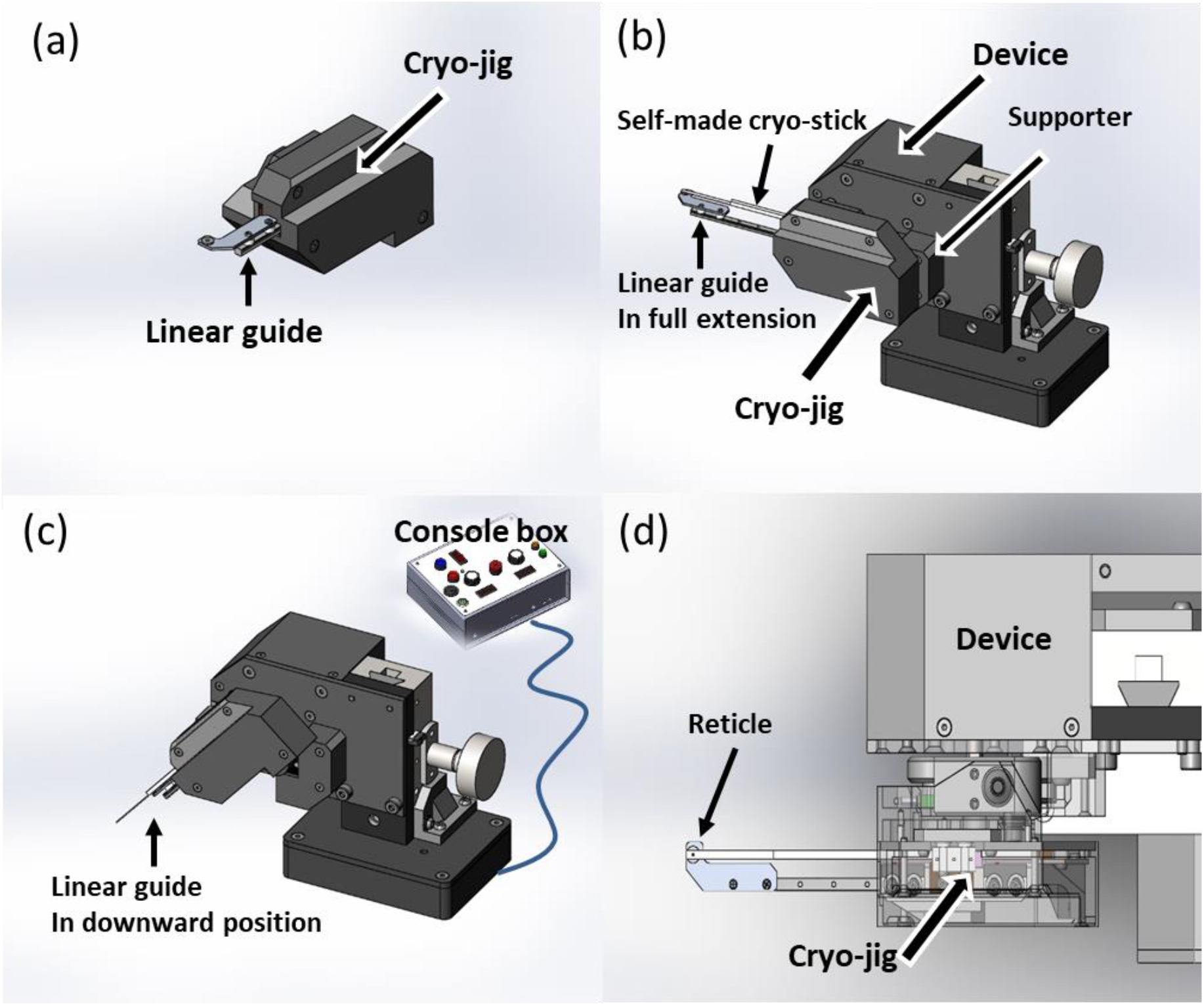
Alignment of Laser machine with the device and cryo-jig. (a) Structure of the cryo-jig with linear guide. (b) The cryo-jig was placed on the supporter of the device with cryo-stick and linear guide in full extension. At this time, the device can be turned on and moved to an appropriate position underneath the laser machine. Then, (c) the console box of the device was operated to swing the jig to the downward position and (d) activated the laser machine to fire the laser beam when the cryo-jig swung back to the upward position. According to the degree of deviation between the position where the laser beam hits the linear guide and the target position, fine adjustments the position of the device was made repeatedly until the laser beam precisely hits the reticle, and then the position of the device was not moved again.

### Cryopreservation

Our customized cryo-jig with self-made cryo-stick was used to rapidly supercool the droplets of VS by plunging them into liquid nitrogen for the long-term cryostorage to allow for equilibration to liquid nitrogen temperature (Fig 2). In the allocated hole of the cryo-jig, the self-made cryostick was inserted as deeply as possible until it reached the stopper. The cryo-jig’s linear guide was extended to its entire length while the mechanism of the cryo-jig automatically maintained its place. The linear guide’s reticle (target point) was positioned at a relative location on the cryojig, and a droplet of the VS was placed onto the cryo-stick blade directly above the reticle. The Cryotop was promptly immersed in liquid nitrogen for long term cryo-storage after being removed from the cryo-jig.

**Fig 2.**
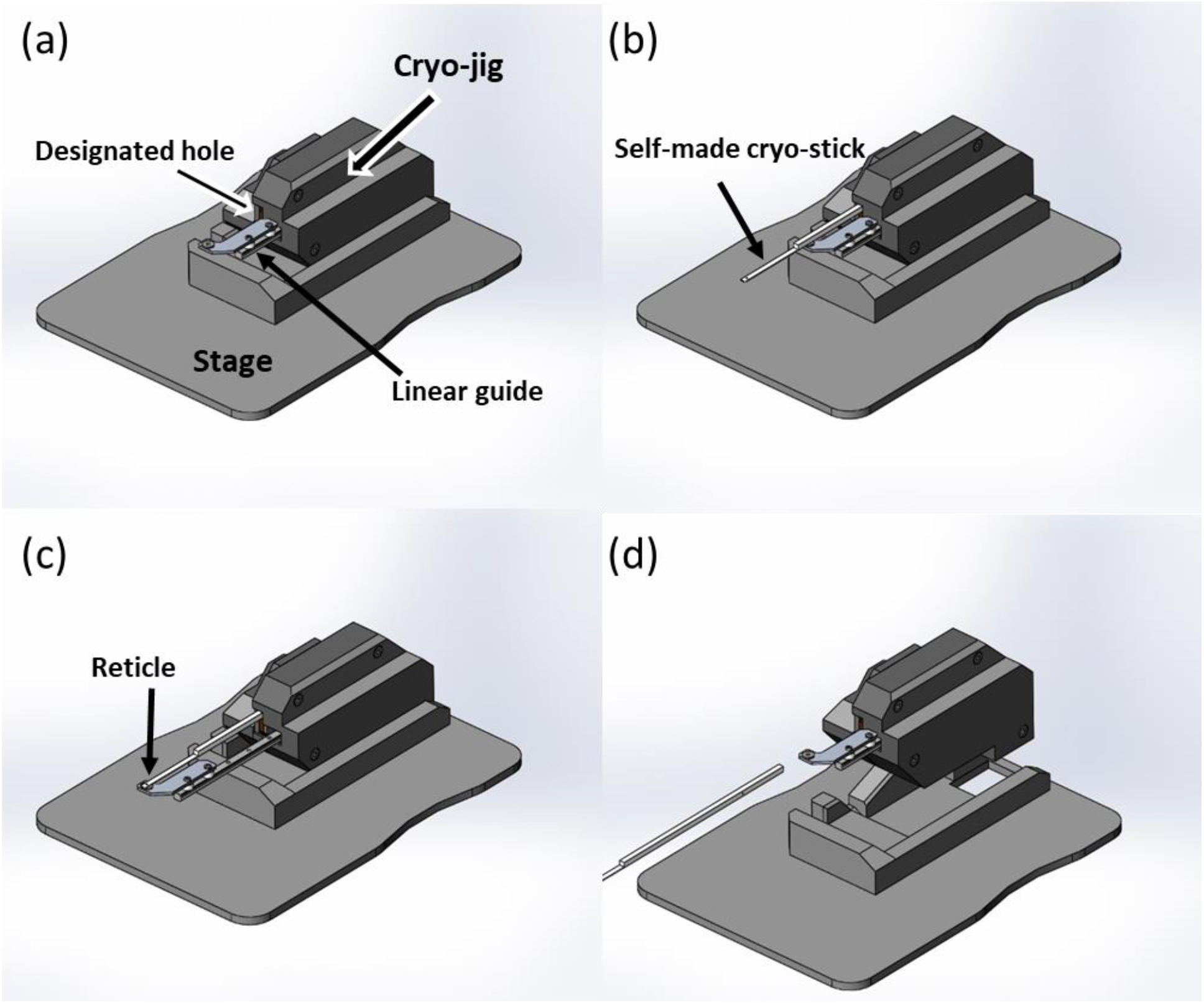
Cryopreservation procedure. (a) The customized cryo-jig with its stage. The designated hole and linear guide were visible on the cryo-jig. (b) The self-made cryostick was inserted to the deepest point until it reached to the stopper in the designated hole of cryo-jig. (c) The linear guide of the jig was pulled out to its full extension while the jig’s mechanism automatically keep its position. The reticle of the linear guide was fixed at a relative position on the jig, and then a drop of about 0.8μl vitrification solution was placed onto the cryo-stick sheet right above the reticle position. (d) The cryo-stick was removed from the cryo-jig for cryo-storage in liquid nitrogen.

### Testing of laser striking accuracy and successful rewarming rate

The laser was used to provide a high energy singular millisecond pulse, subsequently, the droplets of VS was hit by laser beam, and the laser striking accuracy and successful rewarming rates were measured. The laser power was dependent on input voltage and pulse period and provided rapid and uniform warming. The cryo-jig was attached to a laser machine that was constructed of conventional bioplastic polylactic acid (PLA) and an ionic polymer-metal composite before laser warming (Fig 3). Briefly, the cryo-jig was set up and the cryo-stick was prepared for the laser warming process. When the cryo-stick was retrieved, with the sample on the blade part remaining within the liquid nitrogen and the stick part remaining outside, the stick part of the cryo-stick was inserted into the designated hole of the cryo-jig until the deepest stopper was reached. Then the cryo-jig was in a downward position at about 45-degree angle by mounting it onto the device’s supporter, which has its mechanism automatically combined. After the digitized actuator was activated, the device swung the cryo-jig and the cryo-stick from the downward state to the upward position with three different tip travel speeds. The upward position of the cryo-jig was used to raise the sample into the laser beam focal region in order to hit sample, in contrast, the downward position allowed the sample to be placed directly into liquid nitrogen for vitrification. The attached control knobs were used to adjust parameters for appropriate rotary movement of the jig. The digitized actuator drove a central platform that supports the jig, and by turning the knobs ahead of time to set the parameters for the jig with a sample, it enabled a suitable velocity and stopped at the desired position. The robotic arm system of the cryo-jig was implemented from the rotated downward state to the rotated upward position when the actuator was engaged. The laser central console and the laser device both include two safety buttons that govern the laser pulse activation. Both security buttons were the safe design, use and implementation of lasers to minimize the risk of laser accidents. The entire process was recorded by an overhead camera, and the video was used to evaluate if any ice formed inside the sample.

**Fig 3.**
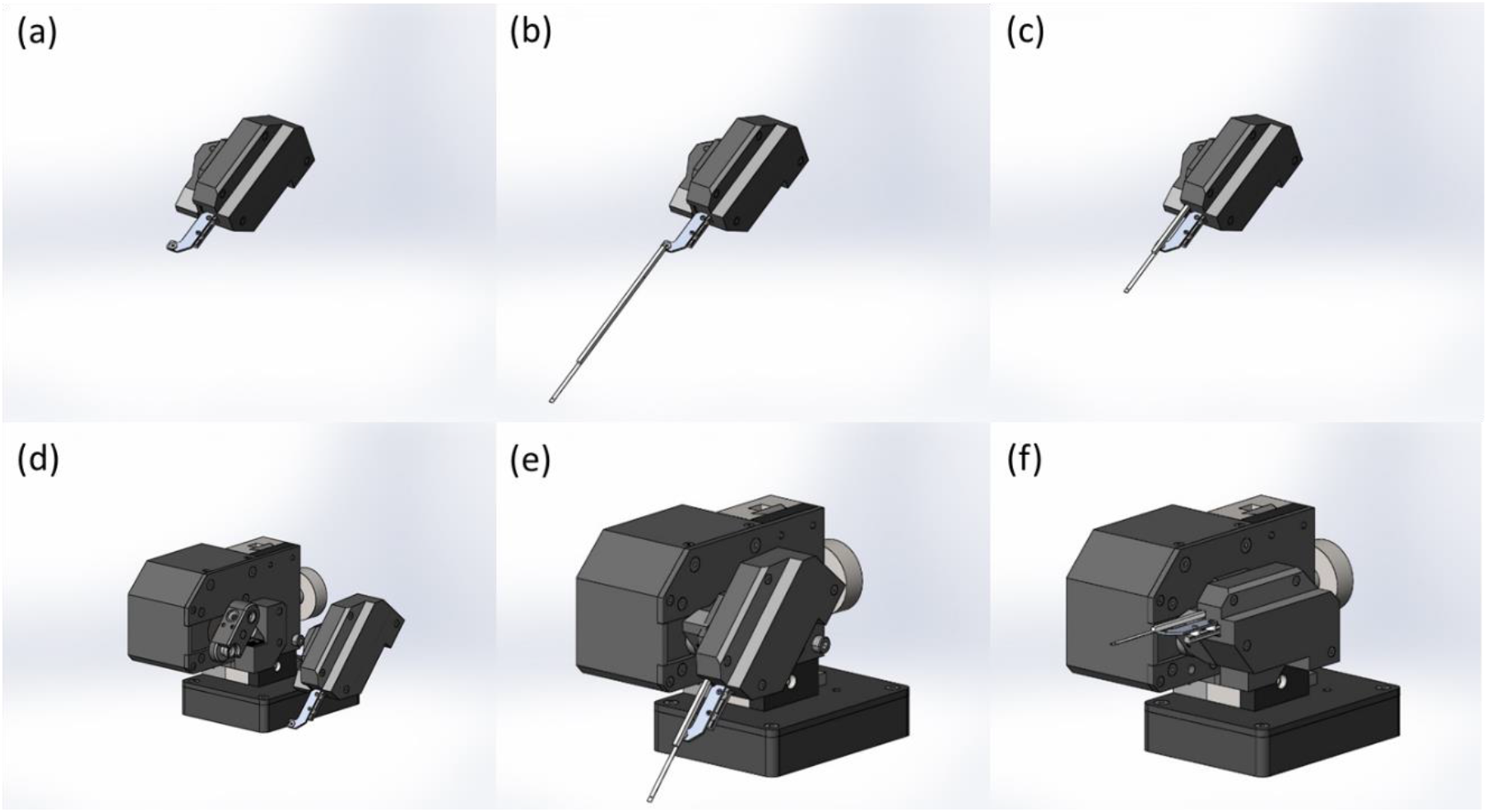
Laser warming procedure. (a) The cryo-jig was in position and ready for laser warming procedure. (b) The sample on the blade of the cryo-stick was kept in liquid nitrogen, while the rest of the cryo-stick was not. (c) The stick part of the cryo-stick was then placed into the designated hole of the cryo-jig until the deepest stopper was reached. (d,e) The cryo-jig was mounted onto the supporter of the device which has its mechanism automatically combined and fixed the cryo-jig to the correct position in a downward state at 45 degree angle. (f) The device swung the cryo-jig with the cryo-stick from the downward to the upward position into the laser beam focus region after the actuator was engaged. When the cryo-stick reached the upper position, the laser hit was instantly activated for warming.

**Figure 4.**
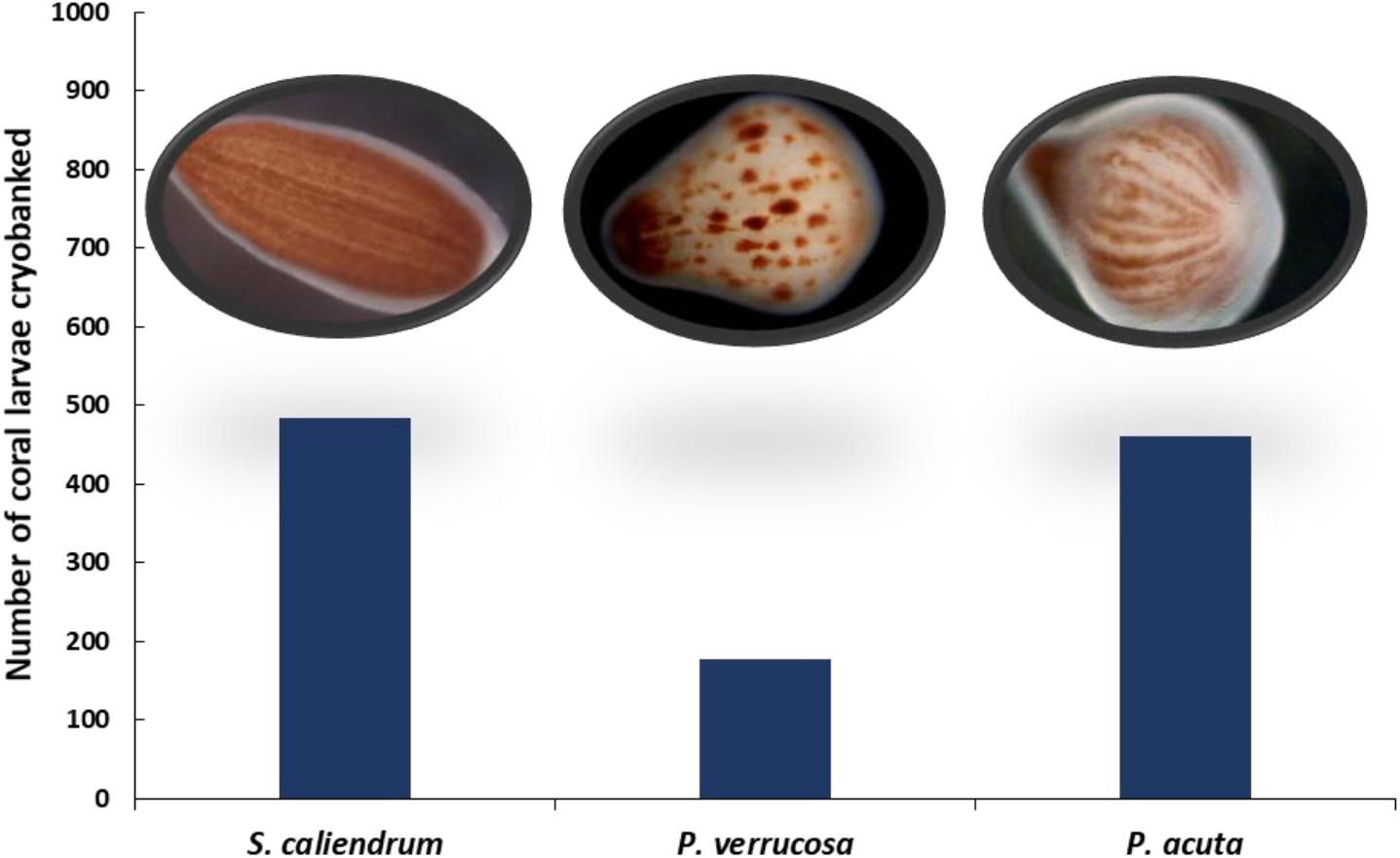
Number of coral larvae *Seriatopora caliendrum, Pocillopora verrucosa* and *P. acuta* cryobanked using the customized device.

Identification of the laser striking accuracy and successful rewarming rate were following the principles below. When the laser was fired, the differences between a laser that hit and one that did not could be seen obviously. Whether the droplet was partially or completely hit could also be differentiated: (1) when the laser hit the droplet, it became crystal clear. If the laser did not hit the droplet, it would turn from translucent to opaque, leaving a light yellow mark on the location where it impacted. (2) When the laser hit the droplet partially, the laser-hitting region would become crystal clear while the non-hitting portion became opaque which would eventually spread to the rest of area (crystallization). On the other hand, when the laser completely hit the droplet, it became crystal clear. The successful rewarming rate was determined when the laser completely hit the droplet, which then became crystal clear without further additional crystallization (opaque) appearance.

### Statistical analysis

Different tip travel speeds were compared using SPSS (version 17.0; SPSS Inc., Chicago, IL, USA) for laser striking accuracy and successful rewarming rate. Using a one-sample Kolmogorov-Smirnov test, the data were examined for normality prior to the analysis of variance. Levene’s test was used to determine if the variance was homogeneous (P > 0.05). The difference was located using one-way analyses of variance (ANOVA) with a Tukey post hoc test. Statistical significance was defined as p values less than 0.05.

## RESULTS AND DISCUSSION

The newly development of cryo-jig and device were able to manipulate the self-made cryo-stick during rapid cooling in liquid nitrogen and ultra-rapid warming by a laser pulse. The sample was super cooled with a vitrification rate of 100% and rapidly warmed by the laser pulse with a high striking accuracy of more than 90%, furthermore, the sample is successfully rewarmed with a highest rate of 62% (Table 1). In this work, we built upon our previous work (Narida et al., 2022) and describe the cryo-jig A and B were operated together to be aligned to hit the same spot, however, combining this device required discretional jig tool (cryo-jig A) and jig holder (cryo-jig B) position and assembling the cryo-stick to reach higher laser striking accuracy and successful rewarming. To address this issue, we focused on developing alignment and bonding techniques for special cryo-jig which were assembling jig tool and jig holder in one piece. The major function of the custom-made work holding cryo-jig is to guide the tool, whereas the fixture is a tool that securely and firmly keeps the work in place to complete a certain operation, and to adjust the manufacturing process in time to increase the laser striking accuracy and consistent vitrification rate, and assure high and successful laser rewarming. This newly produced customized cryo-jig was demonstrated to have significantly higher laser striking accuracy of more than 90% and a successful rewarming rate of 62% in comparison to that presented in our prior study of 76% and a rewarming rate of 59%, respectively (Narida et al., 2022). A higher striking rate would produce more successful rewarming samples. In the present study, the average laser energy represents the mean value of energy per pulse for three different tip travel speed settings that resulted in rewarming of the droplet without any ice crystallization (i.e. going from glassy to cloudy). The droplets were warmed at the highest jig tip travel speed (1113 mm/s), the laser striking accuracy and glassy droplets rate after laser rewarming was 95% and 60% respectively. In contrast, when they were warmed more slow tip travel speed (668 mm/s), it was 92% and55%. Although the difference between these speeds were not statistically significant, it is possible the slower warming speed were consistent with ascribing the lethality to the recrystallization of ice during warming. Further study on formulating higher speeds (i.e., 2000 mm/s or more) from the device to improve successful rewarming is on-going in our lab. With this innovative approach, we can routinely achieve positioning and alignment accuracies of bonded components more efficient and high laser striking accuracy and successful laser rewarming for completing and remaining vitrification state throughout the course of cryopreservation.

**Table 1.** shows the newly designed cryo-jig’s and device average percentage of laser striking accuracy and successful rewarming rate for long-term cryo-storage, vitrification, and laser warming.

| Laser warming (%) |  |  |  |
| --- | --- | --- | --- |
| Tip travel speed | 1113 mm/s | 834 mm/s | 668 mm/s |
| Laser striking accuracy | 95.00 ± 0 | 93.33 ± 2.89 | 91.67 ± 2.89 |
| Successful rewarming rate | 60.00 ± 5.00 | 61.67 ± 2.89 | 55.00 ± 0 |

N=180

Various vitrification carriers are used to manipulate the sample during the cryopreservation processes. Some suggest a loop or hook device that consists of a closed loop or an open hook made of plastic or metal wire attached to the end of a stem and used to pick up biological samples (Lane et al., 1999). Research on cryopreservation of animal cell or tissue were performed using Cryotop for vitrification (Inna et al., 2018) and such Cryotop is normally expensive (each Cryotop costs USD 15-30). In this work, the self-made cryo-stick is a 20 × 0.4 × 0.1 mm in size consisting of a fine blade of transparent film attached to an acrylonitrile butadiene styrene (C8H8.C8H6.C6H6N)N stick backing that is more than ten times less costly. Our newly customized device was specifically designed to be engineered with special made one-piece cryojig. As the present study indicated that the quality of vitrified samples utilizing our self-made cryostick method was evaluated in this work, and it was found that approximately 62% of the samples successful rewarmed and remained vitrified state effectively. The results of our present study strongly suggested that our self-made cryostick coupled with a customized jig and device offered the best ability to produce ultra-rapid cooling and warming with minimal ice crystal formation for this study.

Vitrification prevents the formation of ice crystals inside the cell by the use of combined cryoprotectant and rapid cooling rates to directly transition to the glassy state without crystallization (Seki and Mazur, 2012; Jin et al., 2014; Jin and Mazur. 2015). Although the cryoprotectant concentration could be reduced to less toxic effects on cells while still achieving vitrification, this increases ice nuclei formed during the freezing process and rapid warming is essential to eliminate intracellular ice formation (Tsai et al., 2016). To initiate infrared laser warming, which can heat a sample droplet by up to ~10 ^7^ °C/min, depending on the amount of energy of laser beams delivered to the sample and the quantity of laser absorption of particles present (Jin and Mazur, 2015). A representative run of the system is shown in Figures in which a small aqueous sample (VS consisting of 2 M EG + 1 M PG, 40% w/v Ficoll, and 10% v/v gold nanobars at a final concentration of 1.2 × 10^18^ GNP/m^3^) was initiated by placing a 0.8 µL volume of the droplet on the 0.1 mm blade of a self-made cryostick which was then placed on a self-customized cryojig and device and its blade cooled by immersion in liquid nitrogen. Subsequently laser warming was effected by lifting the cryo-stick blade out of the liquid nitrogen, positioned under the laser, and the laser fired within 0.25s with the characteristic wavelength of 535 nm. For our purposes, the most important fact is that once we had empirically calculated the melting energy (voltage) for a particular droplet size, composition, and gold nano-bar concentration, we could warm repeating droplet preparations in a consistent manner. Notably, we reported on the use of a laser pulse to achieve a great rewarming rate of 62%. The concentration of gold nanoparticles (1.2 × 10^18^ GNP/m^3^) and laser settings (300V, 10ms pulse) used in the present study were selected to produce the minimal amount of warming required to consistently warm the 0.8 µL sample droplet to avoid overheating and to be sufficient to completely prevent the ice recrystallization.

The present study demonstrated significant progress in successful long-term cryostorage of coral larvae of 484 *S. caliendrum*, 176 *P. verrucosa* and 460 *P. acuta* using our newly constructed vitrification device, cryojig and cryo-stick. Cryopreservation of sperm, somatic cells, tissue as well as Symbiodiniaceae isolated from numerous genera was also demonstrated to be effective (Viyakarn et al., 2018; Di Genio et al., 2020; Cirino et al., 2021; Toh et al., 2022; Lin et al., 2022). However, cryo-storage of coral oocytes and embryos is substantially more difficult as they are sensitive to chilling which results in noticeable effect on the cell membrane characteristics, function and integrity (Lin et al., 2014; Tsai et al., 2016; Lin and Tsai, 2020). Major efforts for future cryo-storage of coral oocytes and embryos are currently being established. Herein, the present of study successfully achieved to long-term cryobank of total over a thousand coral larvae, and these biological materials repositories will be a critical safeguard against extinction for corals threatened by climate change, disease, and genetic diversity loss.

## CONCLUSION

This research represented a novel methodology that can be routinely achieved with high alignment precision, robust, ultra-stable optical assemblies with sound performance and stability for long-term cryobanking of various species. We anticipate that as the long-term cryostorage approach was perfected and developed in this study, it will allow for further examples of the application of cryobanking employing vitrification and nano-laser warming to aid a wide range of cells and tissues from diverse species.

## ACKNOWLEDGEMENT

This study was supported by the Ministry of Science and Technology.

## AUTHOR CONTRIBUTIONS

Conceptualization, C.L. and S.T.; methodology, W.C.H., C.L.H., T.C.C and K.L.; validation, C.L. and S.T.; formal analysis, W.C.H., C.L.H., T.C.C and K.L.; resources, C.L. and S.T.; writing— original draft preparation, C.L. and S.T.; writing—review and editing, C.L. and S.T.; visualization, C.L. and S.T.; supervision, C.L. and S.T.; funding acquisition, C.L. All authors have read and agreed to the published version of the manuscript.

**The results related to coral cryobanking and repository described in this preprint will be published separately in details**.

## REFERENCES

Cirino, L., Tsai, S., Wang, L. H., Chen, C. S., Hsieh, W. C., Huang, C. L., Wen, Z. H., & Lin, C. (2021) Supplementation of exogenous lipids via liposomes improves coral larvae settlement post-cryopreservation and nano-laser warming. Cryobiology, 98, 80–86. https://doi.org/10.1016/j.cryobiol.2020.12.004

Cirino, L., Wen, Z. H., Hsieh, K., Huang, C. L., Leong, Q. L., Wang, L. H., Chen, C. S., Daly, J., Tsai, S., & Lin, C. (2019). First instance of settlement by cryopreserved coral larvae in symbiotic association with dinoflagellates. Scientific Reports, 9(1), 18851. https://doi.org/10.1038/s41598-019-55374-6

Di Genio, S., Wang, L.H., Meng, P.J., Tsai, S., Lin, C. (2020). Symbio-Cryobank: towards the development of a cryogenic archive for the coral reef dinoflagellate symbiont Symbiodiniaceae. Biopreserv Biobank 19:91–93. https://doi.org/doi.org/10.1089/bio.2020.0071

Hwang, I. S., Hochi S. (2014) Recent Progress in Cryopreservation of Bovine Oocytes. Biomed Res Int, https://doi.org/10.1155/2014/570647

Inna, N., Sanmee, U., Saeng-anan, U., Piromlertamorn, W., Vutyavanich. T. (2018) Rapid freezing versus Cryotop vitrification of mouse two-cell embryos. Clinical and Experimental Reproductive Medicine, 45(3):110. https://doi.org/10.5653/cerm.2018.45.3.110

Jin, B., & Mazur, P. (2015). High survival of mouse oocytes/embryos after vitrification without permeating cryoprotectants followed by ultra-rapid warming with an IR laser pulse. Scientific Reports, 5, 1–6. https://doi.org/10.1038/srep09271

Jin, B., Kleinhans, F., & Mazur, P. (2014). Survivals of mouse oocytes approach 100% after vitrification in 3-fold diluted media and ultra-rapid warming by an IR laser pulse. Cryobiology, 68(3), 419–430. https://doi.org/10.1016/j.cryobiol.2014.03.005

Khosla, K., Wang, Y., Hagedorn, M., Qin, Z., & Bischof, J. (2017). Gold Nanorod Induced Warming of Embryos from the Cryogenic State Enhances Viability. ACS Nano, 11(8), 7869–7878. https://doi.org/10.1021/acsnano.7b02216

Lane, M., Bavister, B. D., Lyons, E. A., Forest, K. T. (1999) Containerless vitrification of mammalian oocytes and embryos. Nat Biotechnol, 17, 1232–1236. doi:10.1038/70795

Leibo, S. P. (1981) Preservation of ova and embryos by freezing. In: New Technologies in Animal Breeding (Brackett E. G., Scidel G. E. and Seidel S. M. eds) Academic Press, New York, United State of America 127–139.

Lin, C., Kuo, F.W., Chavanich, S., Viyakarn, V. (2014). Membrane lipid phase transition behavior of oocytes from three gorgonian corals in relation to chilling injury. PLoS One 9: e92812. https://doi.org/10.1371/journal.pone.0092812

Lin, C., Tsai, S. (2020). Fifteen years of coral cryopreservation. Platax 17:53–76. https://doi.org/10.29926/platax.202012_17.0004

Lin, C.C., Li, H.H., Tsai, S., Lin, C. (2022). Tissue cryopreservation and cryobanking: Establishment of a cryogenic resource for coral reefs. Biopreserv Biobank 20:409–41. https://doi.org/10.1089/bio.2021.0089

Narida, A., Hsieh, W. C., Huang, C. L., Wen, Z. H., Tsai S., Lin C. (2022) Long-term cryopreservation for vitrification and laser warming. Biopreservation and biobanking https://doi.org/10.1089/bio.2022.0033

Parmegiani, L., Cognigni G.E., Bernardi, S., Cuomo, S., Ciampaglia, W., Infante, F.E., Tabarelli de Fatis, C., Arnone, A., Maccarini, A.M., Filicori, M. (2011) Efficiency of aseptic open vitrification and hermetical cryostorage of human oocytes. Reprod Biomed Online, 23(4), 505–512. https://doi.org/10.1016/j.rbmo.2011.07.003.

Ribeiro, J. C., Carrageta, D. F., Bernardino, R. L., Alves, M. G., Oliveira, P.F. (2022) Aquaporins and Animal Gamete Cryopreservation: Advances and Future Challenges. Animals, 12(3), 359. https://doi.org/10.3390/ani12030359

Seki, S., & Mazur, P. (2012). Ultra-rapid warming yields high survival of mouse oocytes cooled to -196°c in dilutions of a standard vitrification solution. PloS one, 7(4), e36058. https://doi.org/10.1371/journal.pone.0036058

Toh, E.C., Liu, K.L., Tsai, S., Lin, C. (2022). Cryopreservation and cryobanking of cells from 100 coral species. Cells 11:2668. https://doi.org/10.3390/cells11172668

Tsai, S., & Lin, C. (2012). Advantages and applications of cryopreservation in fisheries science. Brazilian Archives of Biology and Technology, 55(3), 425–434. https://doi.org/10.1590/S1516-89132012000300014

Tsai, S., Yang, V., & Lin, C. (2016). Comparison of the cryo-tolerance of vitrified gorgonian oocytes. Scientific Reports, 6, 1–8. https://doi.org/10.1038/srep23290

Tsai, S., Yen, W., Chavanich, S., Viyakarn, V., & Lin, C. (2015). Development of cryopreservation techniques for gorgonian (Junceella juncea) oocytes through vitrification. PLoS one, 10(5), 1–14. https://doi.org/10.1371/journal.pone.0123409

